# Structural insights into the activation of human calcium-sensing receptor

**DOI:** 10.1101/2021.03.30.437720

**Authors:** Xiaochen Chen, Lu Wang, Zhanyu Ding, Qianqian Cui, Li Han, Yongjun Kou, Wenqing Zhang, Haonan Wang, Xiaomin Jia, Mei Dai, Zhenzhong Shi, Yuying Li, Xiyang Li, Yong Geng

## Abstract

Human calcium-sensing receptor (CaSR) is a G-protein-coupled receptor that maintains Ca^2+^ homeostasis in serum. Here, we present the cryo-electron microscopy structures of the CaSR in the inactive and active states. Complemented with previously reported crystal structures of CaSR extracellular domains, it suggests that there are three distinct conformations: inactive, intermediate and active state during the activation. We used a negative allosteric nanobody to stabilize the CaSR in the fully inactive state and found a new binding site for Ca^2+^ ion that acts as a composite agonist with L-amino acid to stabilize the closure of active Venus flytraps. Our data shows that the agonist binding leads to the compaction of the dimer, the proximity of the cysteine-rich domains, the large-scale transitions of 7-transmembrane domains, and the inter-and intrasubunit conformational changes of 7-transmembrane domains to accommodate the downstream transducers. Our results reveal the structural basis for activation mechanisms of the CaSR.

## Introduction

Extracellular calcium ions (Ca^2+^) are required for various kinds of biological processes in the human body (Geng et al., 2016). Human calcium-sensing receptor (CaSR) is a G-protein-coupled receptor (GPCR) that senses small fluctuations of the extracellular levels of Ca^2+^ ions in the blood (Brown et al., 1993). It maintains the Ca^2+^ homeostasis by the modulation of parathyroid hormone (PTH) secretion from parathyroid cells and the regulation of Ca^2+^ reabsorption by the kidney (Brown, 2013). Recently, it has been reported that CaSR is also a phosphate sensor that can sense moderate changes in the extracellular phosphate concentration (Centeno et al., 2019; Chang et al., 2020; Geng et al., 2016). Dysfunctions of CaSR or mutations in its genes may lead to Ca^2+^ homeostatic disorders, such as familial hypocalciuric hypercalcemia, neonatal severe hyperparathyroidism and autosomal dominant hypocalcemia (Hendy et al., 2009; Pollak et al., 1993; Ward et al., 2012).

CaSR belongs to the family C GPCR that includes gamma-aminobutyric acid B (GABA_B_) receptors, metabotropic glutamate receptors (mGluRs), taste receptors, GPCR6a and several orphan receptors (Ellaithy et al., 2020; Hannan et al., 2018; Heaney and Kinney, 2016; Pin and Bettler, 2016). Like most class C GPCRs, CaSR functions as a disulphide-linked homodimer. Each subunit of CaSR is comprised of a large extracellular domains (ECD) that contains a ligand-binding Venus flytrap (VFT) domain and a cysteine rich domain (CRD), and a 7 transmembrane domian (7TMD) that connects to CRD to carry signals from VFT domain to the downstream G proteins (Geng et al., 2016; Zhang et al., 2016).

CaSR can be activated or modulated by Ca^2+^ ions, amino acids (Geng et al., 2016; Zhang et al., 2016), L-1,2,3,4-tetrahydronorharman-3-carboxylic acid (TNCA), a tryptophan derivative ligand (Zhang et al., 2016), and several commercial calcium mimetic drug, such as cinacalcet (Leach et al., 2016; Nemeth et al., 2004), etelcacetide and evocalcet (positive allosteric modulator of the CaSR) that used for patients with end-stage kidney diseases undergoing dialysis (Alexander et al., 2015; Leach et al., 2016; Walter et al., 2013).

The recent groundbreaking structural studies of some class C full-length receptors, such as mGluR5 (Koehl et al., 2019) and GABA_B_ receptors (Kim et al., 2020; Mao et al., 2020; Papasergi-Scott et al., 2020; Park et al., 2020; Shaye et al., 2020) by cryo-electron microscopy (cryo-EM) have provided a structural framework through which to unravel the activation mechanisms of class C GPCRs. The crystal structures of the resting and active conformations of CaSR ECD were solved by two different groups (Geng et al., 2016; Zhang et al., 2016). More recently, the cryo-EM structures of full length CaSR in active and inactive states have been solved (Ling et al., 2021). However, the reported inactive structures do not show the fully inactive state and these results are similar to the previous reported crystal structures of CaSR ECD (Geng et al., 2016). They proposed that Ca^2+^ ions and L-Trp work cooperatively to activate CaSR, which leads to the closure of VFT domain of CaSR, while the proposed mechanisms remain controversial. In our study, we used cryo-EM to obtain the structures of full-length CaSR in inactive and active conformations (Figure 1). The fully inactive structure is stabilized by a negative allosteric nanobody. The active structure has demonstrated a calcium binding site at the interdomain cleft of VFT, and the Ca^2+^ and TNCA constitute of a composite agonist to stabilize the closure of the VFT, leading to the conformational changes of the 7TMDs to initiate signaling.

**Figure 1.**
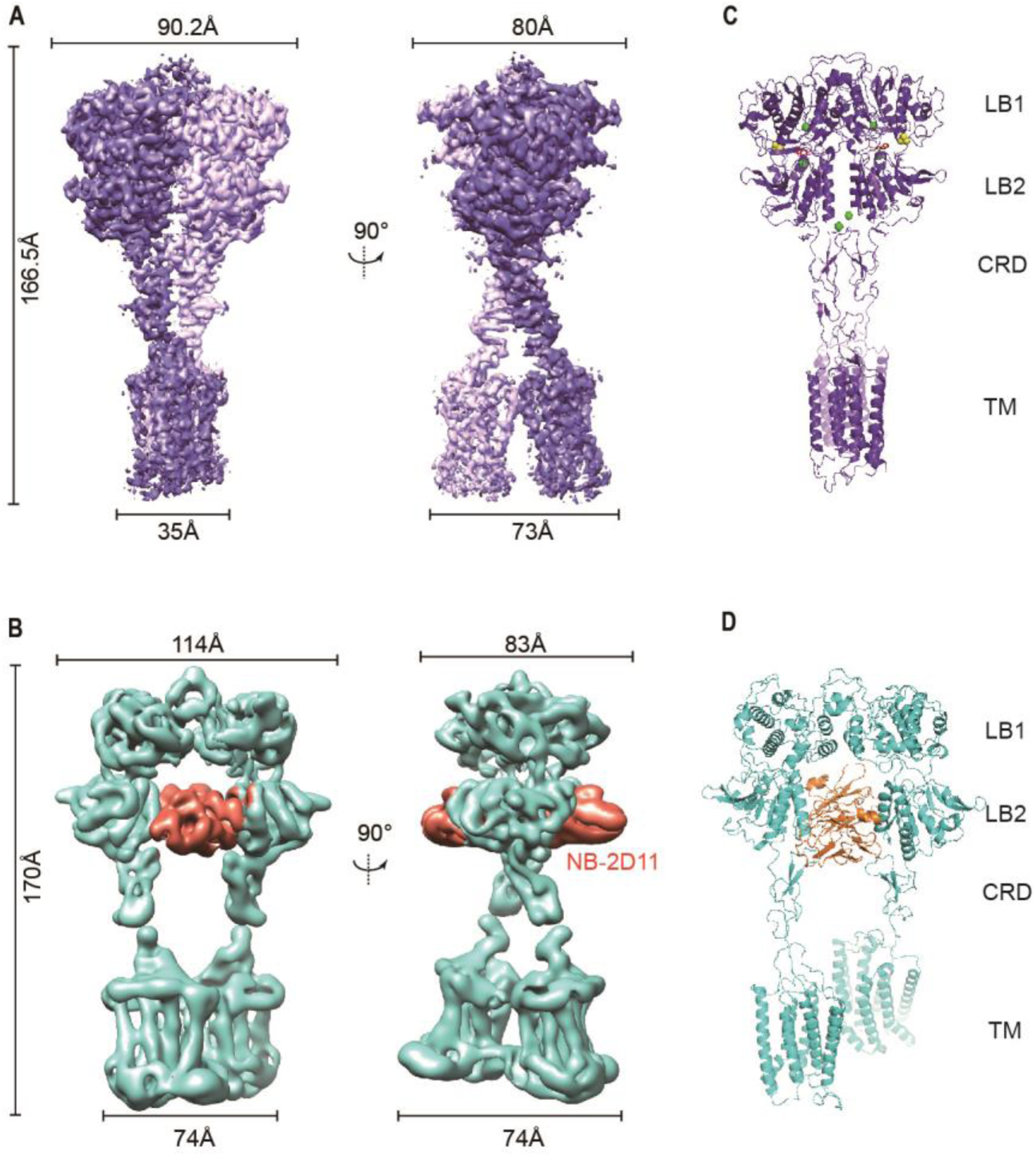
Cryo-EM maps and models of full-length CaSR. (A) Left panel shows the view of CaSR in the active conformation (purple) from front view, and the right panel show the view after a 90° rotation as indicated. (B) Left panel shows the view of CaSR in the inactive conformation (cyan) bound to NB-2D11(orange) from front view, and the right panel show the view after a 90° rotation as indicated. (C) Model (Ribbon representation) of CaSR shows the structure of the active state (purple) bound to TNCA (red) and Ca^2+^ ion (green) viewed from the side. (D) Model (Ribbon representation) of CaSR shows the structure of the inactive state(cyan) bind with NB-2D11 (orange)

## Results

### Determining the cryo-EM structures of full-length CaSR

To obtain the structure of the receptor in active state, we collected a dataset of detergent solubilized full-length CaSR in the presence of positive allosteric modulator (PAM) cinacalcet, 10mM calcium and the compound TNCA. We have observed two active conformations with overall resolutions of 2.99 Å and 3.43Å (Figure 1-figure supplement 1A). We performed local refinement of ECDs and TMDs separately to obtain the maps with the resolutions of 3.07 Å and 4.3 Å, respectively (Figure 1-figure supplement 1A). The high quality density maps have demonstrated well-solved features in the ECD, which allow the unambiguous assignment of calcium, TNCA and most side chains of amino acids of the receptor (Figures 1C; Figure 1-figure supplement 2A). Although the local resolution of 7TMD is low, it enables us to define the backbone of TM helices and even the side chains of some amino acids (Figures 1C; Figure 1-figure supplement 2A). In an attempt to stabilize the structure of the CaSR in the inactive state, we collected a dataset of CaSR in glyco-diosgenin (GDN) formed micelles in the presence of NPS-2143 (a negative allosteric modulator, NAM) and an inhibitory nanobody termed NB-2D11.

The cryo-EM data has demonstrated three conformations of inactive CaSR with an overall resolution of 5.79 Å, 6.88 Å and 7.11 Å, respectively (Figure 1-figure supplement 1B). The local refinement focusing on the ECDs and the 7TMDs was performed separately to improve the resolutions to 4.5 Å and 4.8 Å, respectively (Figure 1-figure supplement 1B), which enabled us to confidently build the backbone of the inactive CaSR model (Figures 1D; Figure 1-figure supplement 2B).

The overall structures in the active and inactive states are homodimeric arrangement that two subunits almost parallelly interact in a side-by-side style while facing opposite directions. For each subunit, the VFT domain is linked to the canonical 7TMD via CRD, which is almost perpendicular to the bilayer membranes (Figure 1B, D). The active structure of CaSR displays a substantial compaction compared to the inactive structure, including the reduction of length, height and width, moreover, their width changed most obviously, because there are four interfaces with interaction between the two promoters at each of LB1 domain, LB2 domain, CRD and 7TM domains (Figure 1) and the closed conformation of the VFT module has shown in the active state, which is stabilized by the TNCA binding at the VFT module and 3 distinct calcium binding sites within each ECD (Figure 1C; Figure 2A). The closure of the VFT is related to TMD through the interaction of the intersubunit CRD. In the inactive structure, there is only one interface at the apex of the receptor and the VFT module adopts an open conformation with the nanobody binding at the left lateral side of each LB2 domain. (Figure 1B, D). The active state has the overall buried surface area of 3378 Å^2^, whereas it substantially decreases to 1346 Å^2^ in the inactive state (Figure 1-figure supplement 2C, D).

**Figure 2.**
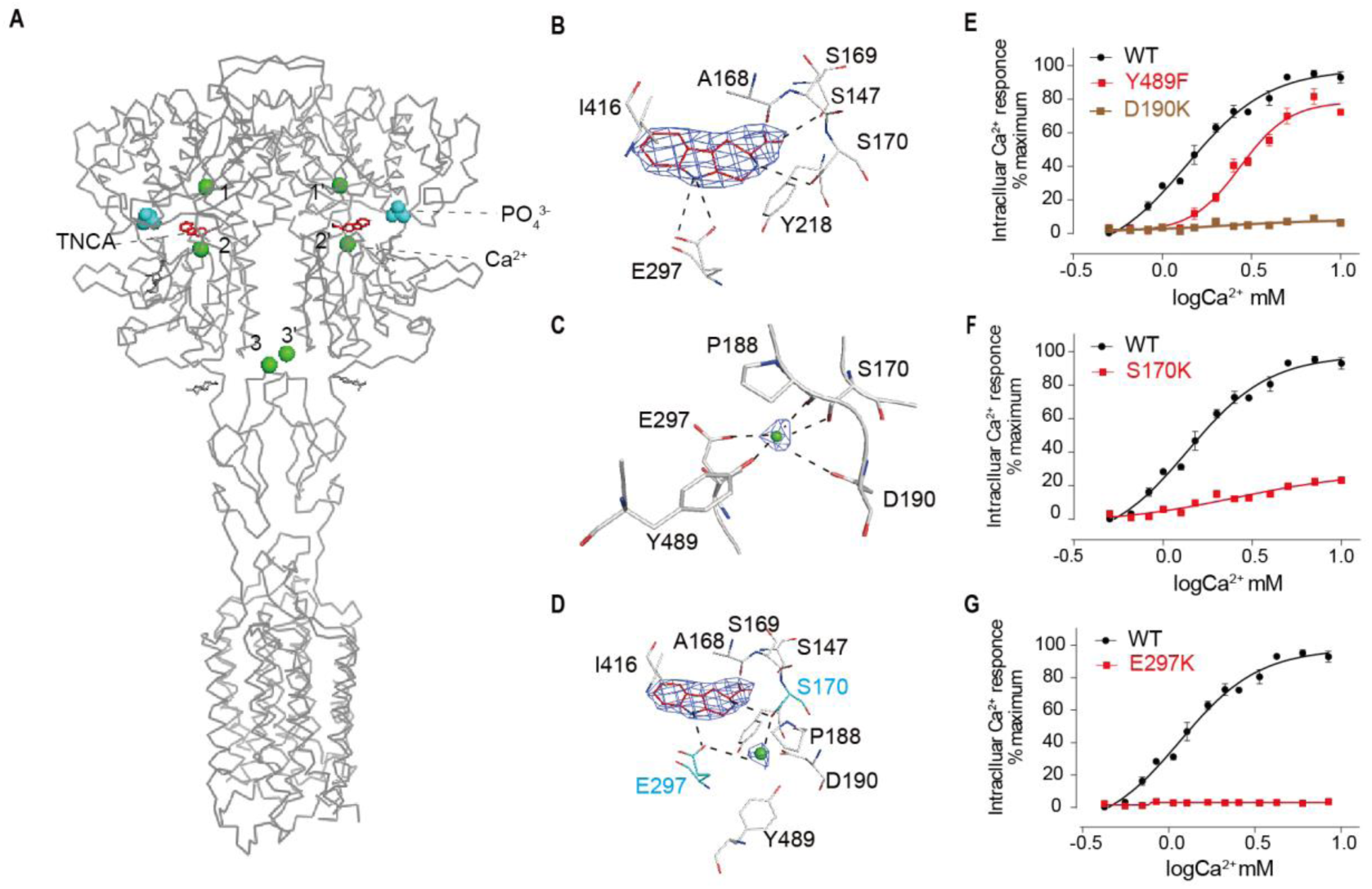
Ca^2+^ and TNCA as a composite against activate the full-length CaSR dimer directly. **(A)** Ribbon representation of the active CaSR (gray), showing the location of the Ca^2+^-binding sites (green sphere) and TNCA (red). **(B)** Specific contacts between CaSR (gray) and TNCA (red space-filling model), mesh represents the final density map contoured at 17σ surrounding. **(C)** Specific interactions between CaSR and newly identified Ca^2+^ ion (green sphere), the mesh represents the cryo-EM density map contoured at 6.0σ surrounding Ca^2+^. **(D)** Highlighting the newly identified Ca^2+^ and TNCA sharing two common binding residues S170 and E297 (cyan space-filling model). **(E)** Dose-dependent intracellular Ca^2+^ mobilization expressing WT (black dots), mutant Y489F (red dots) and mutant D190K (brown dots) CaSR. The single mutations were designed based on Ca^2+^ binding sites. n=4, data present mean ± SEM. **(F-G)** Dose-dependent intracellular Ca^2+^ mobilization expressing WT (black dots), mutant (red dots) CaSR. The single mutations of S170K (F) and 10 E297K (G) were designed based on Ca^2+^ and TNCA binding sites. n=4, data present mean ± SEM.

### Ca^2+^ and TNCA as a composite agonist activate the full-length CaSR dimer directly

The cryo-EM map of active state has shown a distinct density at the ligand-binding cleft of each promoter, which enabled us to unambiguously model TNCA (Figure 2A, B). The binding details of TNCA were the same as the previously reported data (Zhang et al., 2016). The interactions between TNCA and VFT are primarily mediated by the hydrogen bonds (Figure 2B). The high-resolution density of active state map enabled us to identify three distinct Ca^2+^-binding sites within ECD of each promoter (Figure 2A). Two sites were previously reported, while a new Ca^2+^-binding site was found at the interdomain cleft of the VFT module that is close to the hinge loop and abuts the TNCA binding site, and interacts with both LB1 and LB2 domains to facilitate extracellular domain closure (Figure 2A-D). The bound Ca^2+^ ion is primarily coordinated with carbonyl oxygen atoms of P188, D190, S170 and E297, and the hydroxyl group of Y489, among which P188, D190 and S170 are located at LB1 domain, while E297 and Y489 are presented at LB2 domain (Figure 2C, D). The mutations Y489F and D190K significantly reduce the effect of Ca^2+^-stimulated intracellular Ca^2+^ mobilization in cells (Figure 2E). Thus, this Ca^2+^-binding site is very important for stabilizing the closure of VFT.

It is interesting that this bound Ca^2+^ and TNCA share two common binding residues S170 and E297 (Figure 2D). Our experiment has demonstrated that each of single mutations S170A and E297K abolishes Ca^2+^-dependent receptor response (Figure 2F, G), which indicates that the CaSR is synergistically activated by the composite agonist composed of TNCA and Ca^2+^ ions.

### The conformational transition of the LB1 prepares for the ligand binding during the activation of CaSR

Both inactive and active structures reveal that the interface of LB1-LB1 dimer is predominantly a hydrophobic core, which is formed by the residues on two central helices (B and C) of each promoter, including V115, V149, as well as L156 for inactive structure and L112, L156, L159, and F160 for active crystal structure (Figure 3A, B; Figure 3-figure supplement 1A, B). On the dimer interfaces, The B-C helix angle has rotated approximately 28° from inactive state (117°) to active state (89°) (Figure 3A, B). These changes were also reported in mGluR5 from active to apo state with an approximately 59° rotation of the B-C helix angle (Koehl et al., 2019) (Figure 3-figure supplement 1C, D), while this rotation is not observed in the crystal structures of CaSR ECDs, and the inactive crystal structure (PDB: 5K5T) displays the same angle as the one in both active crystal (PDB: 5K5S) and cryo-EM structure (Figure 3B-D). These results have indicated that at least two different conformations of LB1 domains were obtained at the inactive state of CaSR. For ease of description, we designated the two distinct rotation angles of the inactive states as inactive conformation for the cryo-EM inactive form and intermediate conformation for the crystal inactive form.

**Figure 3.**
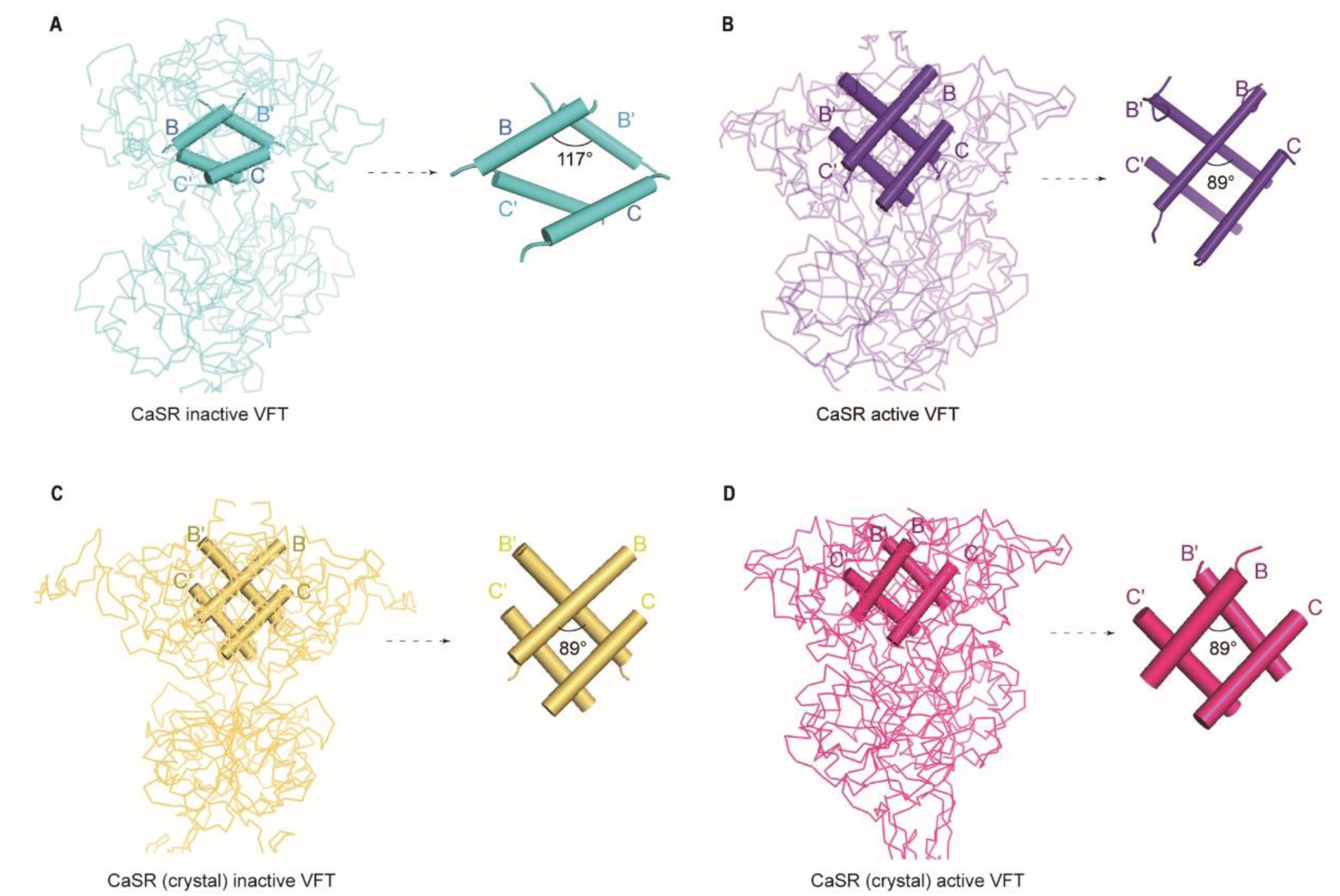
Comparisons of intersubunit LB1 domains interfaces in the inactive and active states of CaSR. **(A)** Left panel: The Cα trace of VFT module of inactive CaSR cryo-EM structure (cyan). Right panel: The B-Helix angle is 117°. **(B)** Left panel: The Cα trace of VFT module of active CaSR cryo-EM structure (purple). Right panel: The B-Helix angle is 89°. **(C)** Left panel: The Cα trace of VFT module of inactive CaSR crystal structure (yellow) (PDB:5K5T). Right panel: The B-Helix angle is 89°. **(D)** Left panel: The Cα trace of VFT module of active CaSR crystal structure (red) (PDB: 5K5S). Right panel: The B-Helix angle is 89°.

The LB1 domain plays a predominant role for anchoring the ligands. Among the superimposition of inactive, intermediate and active conformations, the inactive conformation of the LB1 domain displays a significant change from that of the other two conformations (Figure 3-figure supplement 1E, F), whereas the LB1 domain in the intermediate and the active states is well superimposed with backbone r.m.s.d. of 0.806 Å (Figure 3-figure supplement 1F). Thus, the conformational transition of the LB1 domains from inactive state to intermediate state provide the structural basis for the ligand binding.

### Spontaneous proximity of LB2 domains during the activation

No significant difference of the overall LB2 conformations is observed among the superposition of inactive, intermediate and active structures (Figure 4-figure supplement 1A, B). The cryo-EM structure of CaSR in inactive state displays a relatively large backbone separation distance of 56.26Å between the C-terminal ends of each LB2 domain, while it reduces to 45.8 Å in the intermedium state. A further reduction to 29 Å is observed upon the activation in the active model (Figure 4A-C). Thus, the two LB2 domains gradually approach each other until they interact, whereas this process is not induced by the agonists (Figure 4A-C; Figure 4-figure supplement 1C-E).

**Figure 4.**
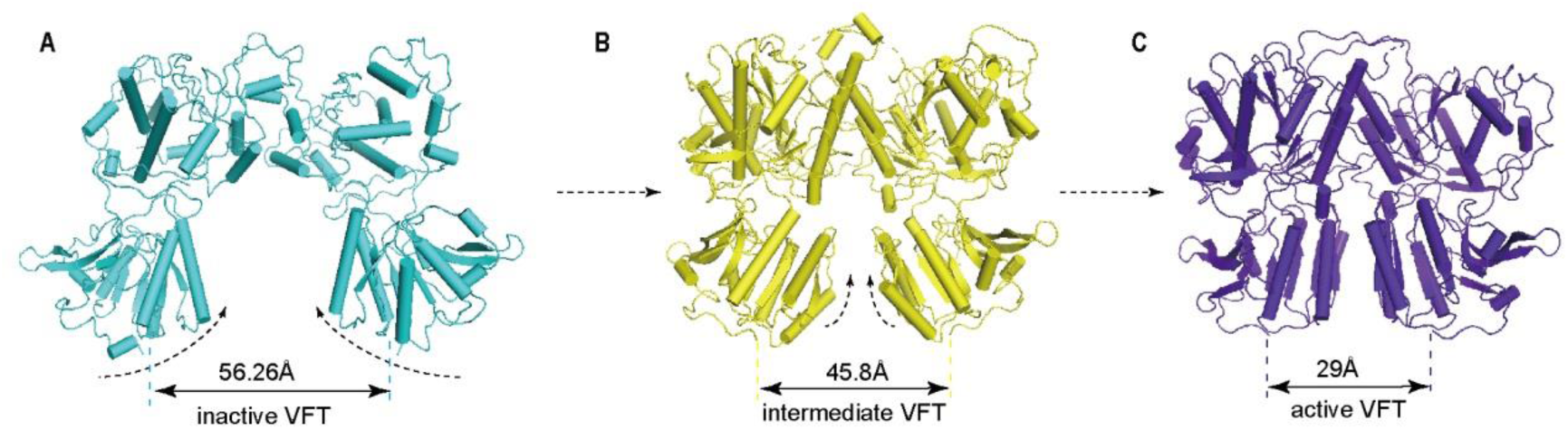
The conformational changes of LB2 domains in three states. **(A)** Inactive (cyan) conformation of VFT module. The distance between C termini of the two LB2 domains is 56.26Å. **(B)** Intermediate (yellow) (PDB:5K5T) conformation of VFT module. The distance between C termini of the two LB2 domains is 45.8Å. **(C)** Active (purple) conformation of VFT module. The distance between C termini of the two LB2 domains is 29Å.

### NB-2D11 blocks the interaction of LB2 domains to trap the CaSR in the inactive conformation

We selected NB-2D11 with a dissociation constant of 0.24 nM (Figure 5A) and IC_50_ value of 41.7 nM (Figure 5B) to stabilize the inactive conformation of CaSR. NB-2D11 binds the left lateral of each LB2 domain from orthogonal view (Figure 5C, D), with the hydrophilic interaction interface between the amino acids D53, D99, W102, R101 and E110 from CDR1 and CDR3 of the nanobody and the residues R220, S240, S244, Y246, S247 and E251 from Helix F and Strand I (Figure 5E). Superposition of the inactive and active LB2 domain shows that NB-D211 occupies the spatial position of the LB2 domain of the other promoter, which blocks the approach of another corresponding subunit LB2 (Figure 5F). Our results indicate that the interactions of both LB2 domains are required to activate CaSR which is the explanation of the inhibitory function of NB-2D11.

**Figure 5.**
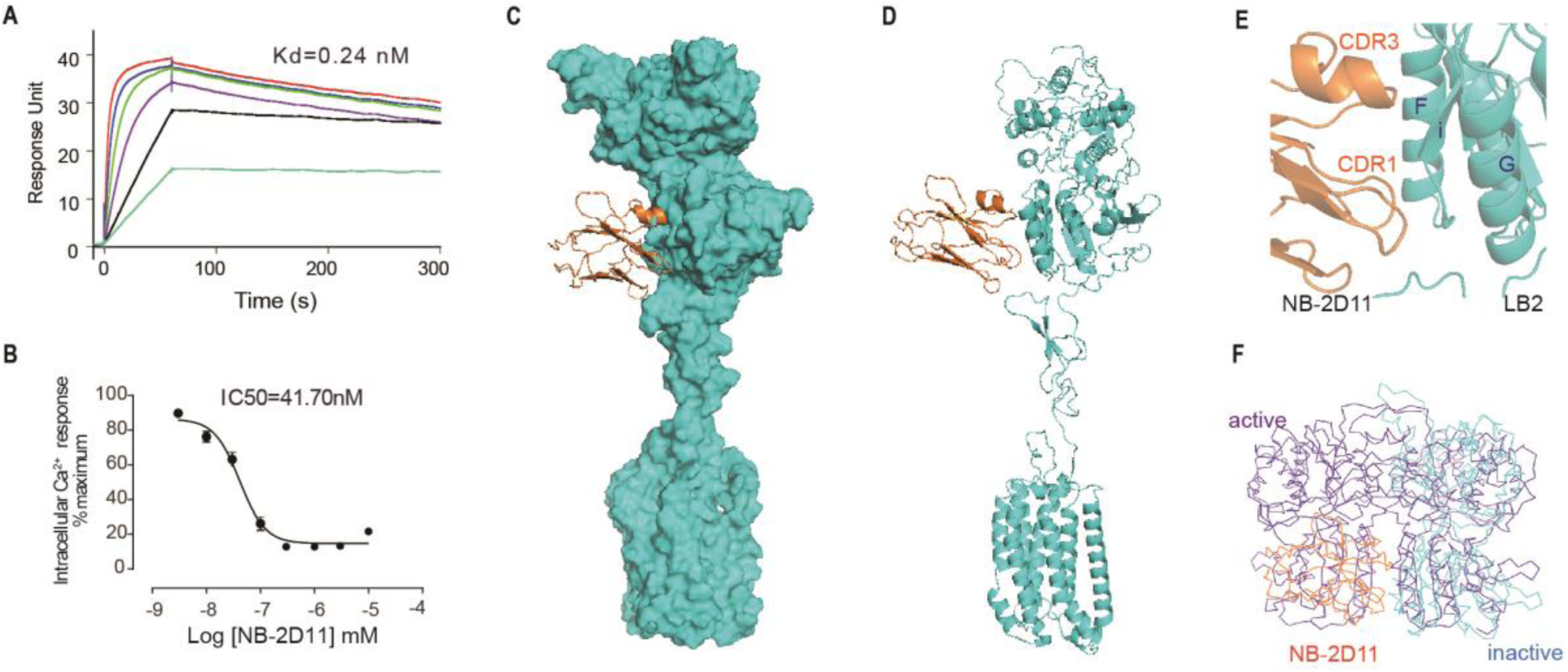
The NB-2D11 blocks the interaction of LB2 domains. **(A)** SPR sensorgram showing that NB-2D11 bound to CaSR with 0.24nM affinity. **(B)** Dose-dependent NB-2D11-inhibited intracellular Ca^2+^ mobilization in response to Ca^2+^ ions. N=3, data present mean ±SEM. **(C)** Structure of the inactive CaSR promoter (surface representation, cyan) with NB-2D11(ribbon diagram, orange) from front view. **(D)** The NB-2D11 (orange) binds the left lateral of the LB2 (cyan) from the front view of the promoter. (**E**) NB-2D11 binds the LB2 domain through a series of polar interaction through CDR1 and CDR3 of the nanobody and the Helix F and Strand I of the CaSR. (**F**) Superposition of NB-2D11(orange) binding inactive conformations (cyan) and active (purple) conformations based on the LB2 domain of VFT module, showing the whole NB-2D11 in the inactive state crashes with the LB2 domain of another VFT module in the active state.

### The rotation of LB2 domain propagates to large-scale transitions of the 7TMDs from TM5-TM6**-**plane to TM6-driven interface

The closure of VFT displays an inward rotation of each LB2 followed by moving upward individually (Figure 4A-C). Afterwards, two intersubunit interfaces are formed at the downstream of subunits, including the interaction between the LB2 linked CRDs, which is consisted with the reported crystal structure of CaSR ECD (Geng et al., 2016) (Figure 6-figure supplement 1) and the intersubunit interaction between TMDs (Figure 6A-F). The alignment of individual 7TMD of both inactive and active states has demonstrated that the helices are well superposed (Figure 6D). Although the NAM and the PAM were added during the preparation of inactive and active samples, respectively, no density of them is observed on the maps due to the low resolutions. The inactive structure reveals that the TM5 and TM6 constitute a 7TMD plane-plane interface (Figure 6E). There are pairwise symmetrical undefined maps that link the extracellular and intracellular part of TM5 and TM6 in the 7TMD interface (Figure 6-figure supplement 2A). Our experiments demonstrated that each of the single mutations F789^5.56^A or F792^5.59^A contacting with the map attenuates Ca^2+^-induced receptor activity, indicating that this contact plays a role in the activation of CaSR (Figure 6H, I). The active structure shows a TM6-TM6 interface, contacting at the apex of TM6 helices which appears to be a sign of activation (Koehl et al., 2019) (Figure 6B, F). To further validate the role of this interface, the mutants of P823R, P823K and A824R markedly reduce the Ca^2+^-induced receptor activity (Figure 6J,K; Figure 6-figure supplement 2B). The previous study showed that a cysteine cross-linking at residue A824^6.56^ led to a constitutively active receptor (Liu et al., 2020). The active mGluR5 (Koehl et al., 2019) and GABA_B_ receptor (Kim et al., 2020; Mao et al., 2020; Papasergi-Scott et al., 2020; Shaye et al., 2020) have the same TM6-TM6 interface as that of CaSR (Figure 6F; Figure 6-figure supplement 2C, D), and it was observed that the TM6 cross-linked mGluR5 and TM6-locked mGluR2 were activated continuously (Koehl et al., 2019; Xue et al., 2015).

**Figure 6.**
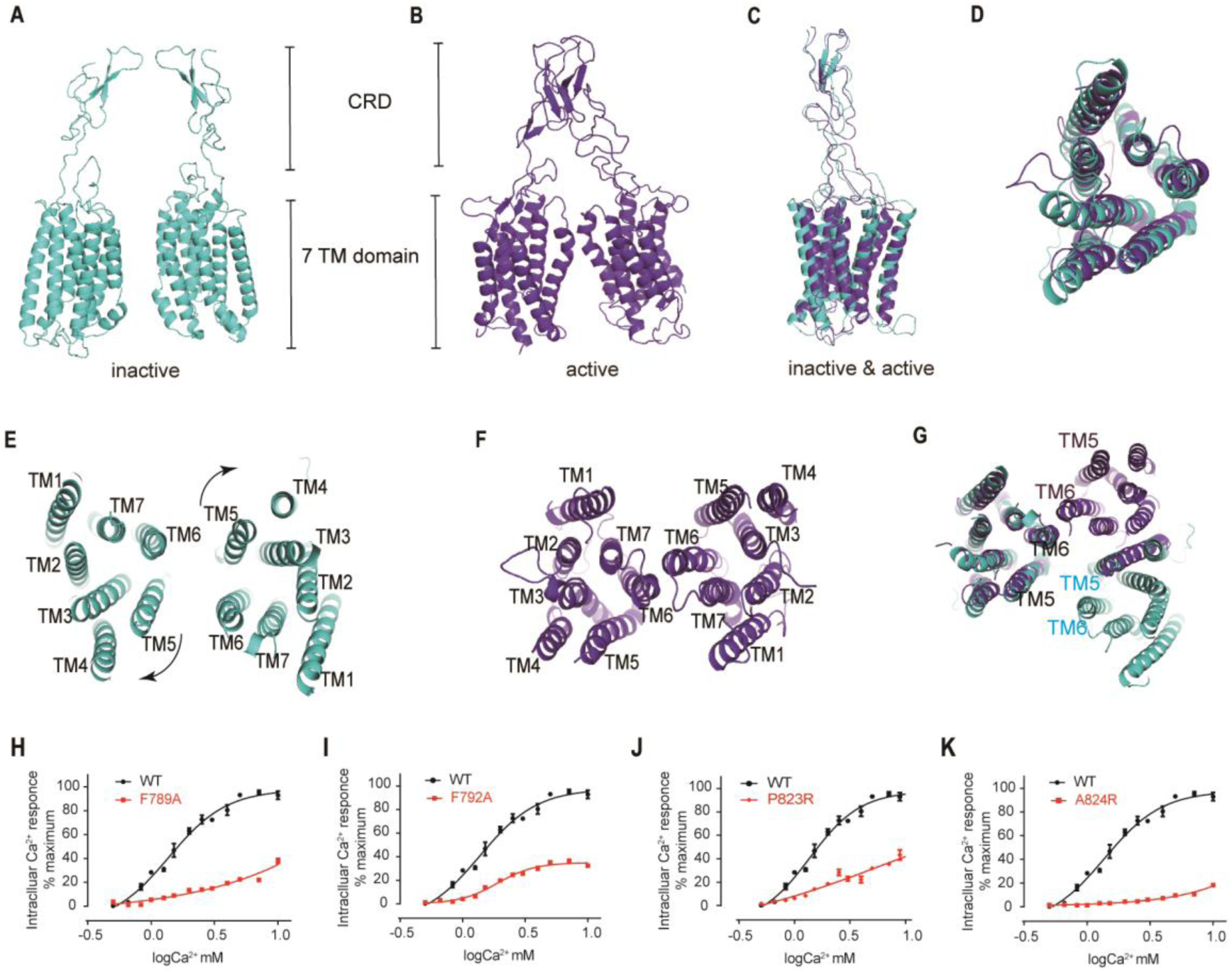
The closure of VFT leading to the rearrangement of inter-7TMDs. **(A)** Front view of inactive CaSR CRDs and 7TMDs (cyan). **(B)** Front view of active CaSR CRDs and 7TMDs (purple). **(C)** The alignment of the part of CRD and 7TMDs in both inactive and active CaSR. **(D)** The alignment of inactive and active 7TMDs from top view. **(E-F)** The rotations of LB2 domains rearrange the 7TMD interface from the interface mediated by TM5-TM6 in the inactive state (E) to the TM6 contacting interface in the active state (F). **(G)**The 7TMDs interface in the inactive state of CaSR is mediated by TM5 and TM6 (cyan) from top view, and that of the active state is driven by TM6 from top view. Superposition of 7TMD of the inactive (cyan) and active CaSR (purple) show the rotation of 7TMDs. **(H-K)** Dose-dependent intracellular Ca^2+^ mobilization expressing WT (black dots) and mutant (red dots) CaSR. The single mutations of F789A (H), F792A (I), P823R (J) and P824R (K) were designed based on the inactive density map. For (H to K), n=4, data present mean ± SEM.

The conformation of the downstream portion of each subunit from the C term of the LB2 domain in both inactive and active states presents similar, except that the bundle composed of ECL2 and the stalk linking CRD and TM1 has little fluctuation, it seems that overall part is semi rigid (Figure 6C). Therefore, a little rotation of LB2 domains could propagate to large-scale transitions of the TMDs through the CRDs, which reorients the 7TMDs from the inactive plane-plane interface mediated by TM5 and TM6 to the active interface driven by TM6 (Figure 6E-G). The proximity of 7TMDs is observed during the activation, from a plane-plane distance of 24 Å in inactive state to 5.7 Å at P823^6.55^ in the active state (Figure 1-figure supplement 2A, B).

### Upward movement of LB2 converted into the intra-7TM rearrangement through ECL2

Both inactive and active structures show that there is a bundle structure in the junction region between extracellular and transmembrane domain, which is composed of C-terminal elongated peptide of CRD and the twisted hairpin loop of ECL2 (Figure 7A, B). Unlike mGluR5 and GABA_B_ receptor (Kim et al., 2020; Koehl et al., 2019; Mao et al., 2020; Papasergi-Scott et al., 2020; Park et al., 2020; Shaye et al., 2020), which formed by a twisted three-strand β-sheet, the junction of CaSR is more flexible than that of mGluR5 and GABA_B_ receptor. There is a common interface constitute of residues 759-763 at ECL2 and residues 601-604 at the C-terminal of CRD (Figure 7A), and the interface in the active conformation is more compact than that of in the inactive conformation (Figure 7B). In addition, there is another interface involved in the residues E759 at the apical loop of ECL2 and the residues W590 at the bottom of the loop composed of residues 589-591 for active state. In the active state, the loop of ECL2 is pulled up by the interaction among E759, W590 and K601, leading to the movement of ECL2 (Figure 7B, C), which was thought to cause the reorientations of TM4 and TM5 domains during the activation of CaSR (Figure 7C). The deletion of residues D758 and E759 at the apex of ECL2 significantly reduces Ca^2+^-induced receptor activity (Figure 7D). The mutations of K601E and W590E disrupting these contact leads to a largely decreased effect of Ca^2+^-stimulated intracellular Ca^2+^ mobilization in cells (Figure 7E, F). Therefore, ECL2 relay the conformational changes of VFT to the intrasubunit TM domain to rearrange the structure to adapt to downstream transducers such as G proteins.

**Figure 7.**
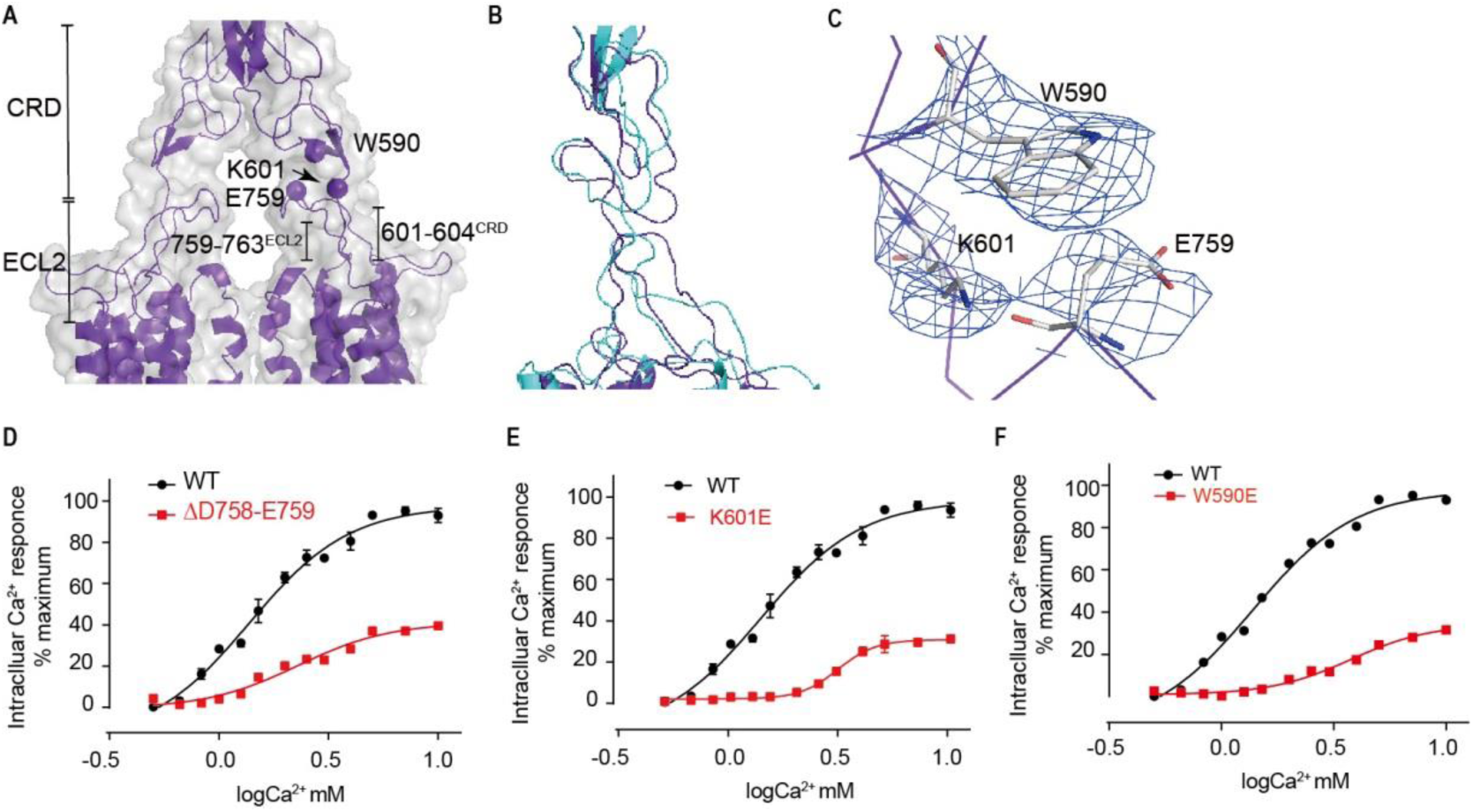
Upward movement of LB2 is converted into the intra-7TM conformational rearrangement through ECL2. **(A)** Model in active state (purple) and cryo-EM map (grey) showing the contact between the CRD and the ECL2 of the 7TMDs. Critical residues at this interface are shown as spheres at their 5 Cα positions. **(B)** Superposition of the interface between the CRD and the ECL2 of the 7TMD between both inactive and active configurations. **(C)** Specific contacts between the loop of CR domain and the loop of ECL2 to shift the ECL2 up. **(D-F)** Deletion of residues D758 and E759 (D), the single mutation of K601E (E) and W590E (F) significantly reduced Ca^2+^-induced receptor activity. (WT in black dots and mutant in red dots). For (D-F), n=4, data present mean ± SEM.

## Discussion

In this study, we have determined the cryo-EM structures of the physiologically and pharmacologically CaSR in inactive and active conformations. The combination of the present two structures of the full-length CaSR, the reported crystal structures of CaSR ECDs and our other biochemical data allows us to understand the structural framework and some essential events for the activation of CaSR. It suggests that there are three distinct conformations in the process of CaSR activation: inactive, intermediate and active state. The overall structures of CaSR resemble the recently published structures of mGluR5 and GABA_B_ receptor (Kim et al., 2020; Koehl et al., 2019; Mao et al., 2020; Papasergi-Scott et al., 2020; Park et al., 2020; Shaye et al., 2020). These three states are automatically converted.

However, in class C GPCR, CaSR has some unique characteristics. The most remarkable feature is that the functions of CaSR can be regulated by Ca^2+^ ions and L-amino acids, while others are activated by L-amino acids or their analogs (Conigrave et al., 2004). There were always the contradictory opinions which the Ca^2+^ is the allosteric activators or agonist or co-agonist with L-amino acids for CaSR. We have identified a new Ca^2+^ ion binding site at the interdomain cleft of the VFT module in the active state. This Ca^2+^ ion shares some common binding residues with TNCA. Therefore, we conclude that the bound Ca^2+^ ion cooperates with TNCA as a composite agonist of CaSR to jointly stabilize the active state.

We try to decipher the occurrence of three important events before the ligand and Ca^2+^ ion. Firstly, the rotation of LB1-LB1 domains is a watershed between inactive and intermediate state. Secondly, the configuration of LB1 domain in the intermediate state is ready for agonist and Ca^2+^ binding, and thirdly, the closure of VFT that stabilized by the agonist and Ca^2+^ ion is a landmark event during the CaSR activation. Previous FRET study has demonstrated the huge conformational changes of CaSR in LB1 domains upon the activation (Liu et al., 2020). Increased glutamate affinity and occupancy in mGluR2 active conformation were observed by the mutations in B-Helix (Levitz et al., 2016). Together with our findings, conformational transitions of the LB1 domains facilitate the ligand binding and prepare for the activation of the CaSR.

It is observed that the LB2 domains approach each other during the activation. Although we do not have a fully active conformation of the CaSR without agonist binding as evidence, the crystal structures of a fully closed VFT modules of mGluR1 with or without agonist binding were previously reported (Kunishima et al., 2000), which indicates that the proximity of both LB2 domains is an automatic process rather than an agonist-driven one. Moreover, the receptor was inhibited if this process was disrupted by the NB-2D11.

We analyzed how the closure of ligand bond VFT module is relayed to the signaling 7TMDs through the CRDs. First, the rotation of LB2 domain is propagated to the large-scale transition of intersubunit 7TMDs, which leads to rearrangement of 7TMDs interface from TM5-TM6-plane/TM5-TM6-plane interface to TM6-TM6 mediated interface that this contact is considered to be a hallmark of activation in class C GPCR. For the mGluR5 and GABA_B_ receptor, the 7TMDs rearrange from TM5-TM5 interface in the inactive state to TM6-TM6 interface in the active state (Kim et al., 2020; Koehl et al., 2019; Mao et al., 2020; Papasergi-Scott et al., 2020; Park et al., 2020; Shaye et al., 2020). The structure provides important insights for drug design if the TM5-TM6-plane interface of 7TMD is unique for CaSR. In addition, the undefined maps that between the 7TMD interface could be some sterols to separate the dimer plane-plane interface and stabilize the inactive state (Figure 6-figure supplement 2A), as the structures of GABA_B_ receptor revealed some cholesterol molecules at the interface of 7TMDs (Kim et al., 2020; Mao et al., 2020; Papasergi-Scott et al., 2020; Park et al., 2020; Shaye et al., 2020).

In this study, it is found that the conformational of ECL2 changed from the inactive to active state. however, the alignment of individual 7TM of both inactive and active-like states has demonstrated that the helices are well superposed, indicating that it is unable to drive the rearrangements of TM4 and TM5 helices and stabilize the active conformation in the TMDs (Figure 6D). This observation is consisted with the reported mGluR5 as the active state be generally stabilized by G proteins (Koehl et al., 2019; Manglik et al., 2015; Rosenbaum et al., 2011). It is required to determine the structure of G protein coupling CaSR to clarify the configuration of the 7TMD in the active state.

## Materials and Methods

### Nanobody library generation

Camel immunizations and nanobody library generation were performed as described previously (Pardon et al., 2014). The animal work is under the supervision of Shanghai Institute of Materia Medica, Chinese Academy of Sciences. In brief, two camels were immunized subcutaneously with approximately 1mg human CaSR protein combined with equal volume of Gerbu FAMA adjuvant once a week for 7 consecutive weeks. Three days after the last immunization, the peripheral blood lymphocytes (PBLs) were isolated from the whole blood using Ficoll-Paque Plus according to manufacturer’s instructions. Total RNA from the PBLs was extracted and reverse transcribed into cDNA using a Super-Script^TM^ III FIRST-Strand SUPERMIX Kit (Invitrogen). The VHH encoding sequences were amplified with two-step enriched-nested PCR using VHH-specific primers and cloned between *PstI* and *BsteII* sites of pMECS vector. Electro-competent *E.coli* TG1 cells were transformed and the size of the constructed nanobody library was evaluated by counting the number of bacterial colonies. Colonies were harvested and stored at −80°C.

### Nanobody identification by phage display

*E.coli* TG1 cells containing the VHH library were superinfected with M13KO7 helper phages to obtain a library of VHH-presenting phages. Phages presenting CaSR-specific VHHs were enriched after 3 rounds of biopanning. For each panning round, phages were dispensed into CaSR coated 96 wells (F96 Maxisorp, Nuc). and incubated for 2 hours on a vibrating platform (700 r.p.m), and subsequently washed 10 times with PBST and 5 times with PBS. The retained phages were eluted with 0.25 mg ml^-1^ trypsin (Sigma-Aldrich). The collected phages were subsequently amplified in *E. coli* TG1 cell for consecutive rounds of panning. After the third round of biopanning, 200 positive clones were picked and infected with M13KO7 helper phages to obtain the VHH-presenting phages.

### Enzyme-linked immunosorbent assays (ELISA) to select CaSR VHHs

The wells of ELISA plates were coated with 2μg ml^-1^ neutravidin in PBS overnight at 4°C. 2μg ml^-1^ biotinylated CaSR was added into each well. Then the wells were blocked by 5mg ml^-1^ non-fat milk powder in PBS. 100μl supernatant of HA-tagged CaSR VHH was added into each well with 1 hour incubation at 4°C, followed by incubation with horseradish peroxidase (HRP)-conjugated anti-HA (Yeasen). TMB substrate (Thermo Fisher Scientific) was added and the reactions were stopped by 2M H_2_SO_4_. The measurement was performed at 450nm.

### Purification of NB-2D11

NB-2D11 was cloned into a pMECS vector that contains a PelB signal peptide and a haemagglutinin (HA) tag followed by a 6×histidine tag at the C-terminus. It was expressed in the periplasm of *Escherichia coli* strain TOP10F’ and grown to a density at OD_600nm_ of 0.6-0.8 at 37℃ in 2YT media containing 100 μg/mL Ampicillin, 0.1% (w/v) glucose and 1mM MgCl_2_, and then induced with 1mM IPTG at 28℃ for 12 hours. The bacteria were harvested by centrifugation and resuspended in the buffer containing 20 mM HEPES pH 7.5, 150 mM NaCl, 1 mM PMSF, and lysed by sonication, then centrifuged at 4,000 r.p.m. to remove cell debris. The supernatant was loaded onto Ni-NTA resin and further eluted in elution buffer containing 20 mM HEPES pH 7.5, 150 mM NaCl, and 300 mM imidazole. The elution was purified by gel filtration chromatography using a HiLoad 16/600 Superdex 75 pg column in 150mM NaCl with 20mM HEPES pH7.5. Finally, NB-2D11 was flash frozen in liquid nitrogen until further use.

### Purification of inactive state CaSR bound to NPS-2143 and NB-2D11

Human CaSR (1-870) followed by a Flag epitope tag (DYKDDDD) at the C-terminus was cloned into a modified pEG BacMam vector (Goehring et al., 2014) for expression in baculovirus-infected mammalian cells. Human embryonic kidney (HEK) 293 GnTI^-^ cells were infected with baculovirus at a density of 2.5×10^6^ cells per ml at 37℃ in 8% CO_2_. 10mM sodium butyrate was added 12-16h post infection, then cells grown for 48h at 30℃ with gentle rotation.

The infected cells were harvested by centrifugation at 4000g for 30 min and resuspended and homogenized using a dounce tissue grinder (WHEATON) in hypotonic buffer (20mM HEPES pH7.5, 10mM NaCl, 1mM CaCl_2_, 10% glycerol, 1×cocktail of protease inhibitor and 1μM NPS-2143). Cell membrane was collected by ultra-centrifugation at 40,000 r.p.m. in a Ti-45 rotor (Beckman Coulter) for 1h. Then the membrane was resuspended and solubilized in buffer containing 20mM HEPES, 150mM NaCl, 1mM CaCl_2_, 10% glycerol, 1μM NPS-2143, 1% (w/v) lauryl maltose neopentyl glycol (LMNG) (Anatrace) and 0.1% (w/v) cholesteryl hemisuccinate TRIS salt (CHS) (Anatrace) for 1 h at 4℃ with constant stirring. The supernatant was collected by ultra-centrifugation at 40,000 r.p.m. for 1h, and applied to an anti-Flag M2 antibody affinity column (Sigma-Aldrich). After receptor binding to the M2 column, the resin was washed with 20mM HEPES, 150mM NaCl, 1mM CaCl_2_, 10% glycerol, 1μM NPS-2143, 0.1% LMNG, 0.01% CHS. The column was washed stepwise with decreasing proportion of LMNG and increasing concentration of GDN/CHS to 0.2%/0.02%. CaSR was then eluted with 20mM HEPES, 150mM NaCl, 1mM CaCl_2_, 10% glycerol, 1μM NPS-2143, 0.02% GDN, 0.002% CHS and 0.2 mg ml^-1^ Flag peptide.

CaSR was further purified by ion-exchange chromatography using a Mono Q 5/50 GL column. Peak fractions were assembled and incubated with a 1.2 molar excess of NB-2D11 for 1h before injection on a Superose 6 increase 10/300 GL column. The fractions of CaSR-NB-2D11 complex in buffer containing 20mM HEPES, 150mM NaCl, 1mM CaCl_2_, 1μM NPS-2143, 0.002% GDN, 0.0002% CHS were pooled and concentrated to approximately 5mg ml^-1^ for further cryo-EM samples preparation.

### Purification of active state CaSR bound to cinacalcet and TNCA

Infected cells (described above) were collected and resuspended in hypotonic buffer (20mM HEPES pH7.5, 10mM NaCl, 10mM CaCl_2_, 10% glycerol, 1×cocktail of protease inhibitor, 1μM cinacalcet and 1μM TNCA). Cell membrane was collected by ultra-centrifugation at 40,000 r.p.m. for 1 h followed by resuspended and solubilized in buffer containing 20mM HEPES, 150mM NaCl, 10mM CaCl_2_, 10% glycerol, 1μM cinacalcet, 1μM TNCA, 1% LMNG and 0.1% CHS for 1h at 4℃. The supernatant was collected by ultra-centrifugation and applied to an anti-Flag M2 antibody affinity column. After receptor binding to the M2 column, the resin was washed with 20mM HEPES, 150mM NaCl, 10mM CaCl_2_, 10% glycerol, 1μM cinacalcet, 1μM TNCA, 0.1% LMNG, 0.01% CHS. LMNG was exchanged for GDN to a proportion of 0.2% in stepwise. CaSR was then eluted with 20mM HEPES, 150mM NaCl, 10mM CaCl_2_, 10% glycerol, 1μM cinacalcet, 1μM TNCA, 0.02% GDN, 0.002% CHS and 0.2 mg ml^-1^ Flag peptide.

CaSR was further purified by Mono Q 5/50 GL column. Peak fractions were assembled and injected to a Superose 6 increase 10/300 GL column. The fractions of CaSR in buffer containing 20mM HEPES, 150mM NaCl, 10mM CaCl_2_, 1μM cinacalcet, 1μM TNCA, 0.002% GDN, 0.0002% CHS were pooled and concentrated to approximately 5mg ml^-1^ for further cryo-EM samples preparation.

### Cryo-EM sample preparation and data acquisition

3μL of inactive or active CaSR protein was applied to glow-discharged holey carbon 300 mesh grids (Quantifoil Au R12/1.3, Quantifoil MicroTools), respectively. The grids were blotted for 2s and flash-frozen in liquid ethane using a Vitrobot Mark IV (Thermo Fisher Scientific) at 4℃ and 100% humidity. Cryo-EM data was collected on a Titan Krios microscope (Thermo Fisher Scientific) at 300kV accelerating voltage equipped with a Gatan K3 Summit direct election detector at a nominal magnification of 81,000× in counting mode at a pixel size of 1.071 Å. Each micrograph contains 36 movie frames with a total accumulated dose of 70 electrons per Å. The defocus range was set -1.5 to -2.5 μm. A total of 5,706 and 4,981 movies for active and inactive CaSR were collected for further data processing, respectively.

### Data processing and 3D reconstruction

All images were aligned and summed using MotionCor2 (Zheng et al., 2017). Unless otherwise specified, single-particle analysis was mainly executed in RELION 3.1 (Zivanov et al., 2020). After CTF parameter determination using CTFFIND4 (Rohou and Grigorieff, 2015), particle auto-picking, manual particle checking, and reference-free 2D classification, 1,546,992 and 2,208,402 particles remained in the active and inactive datasets, respectively. The particles were extracted on a binned dataset with a pixel size of 4.42 Å and subjected for 3D classification, with the initial model generated by ab-initio reconstruction in cryoSPARC (Punjani et al., 2017).

For the CaSR active state dataset, the 3D classification resulted in extraction of 36.6% good particles with a pixel size of 1.071 Å. The particles were subsequently subjected to an auto-refine procedure, yielding a 4.3-Å-resolution map. Afterwards, particles were polished, sorted by carrying out multiple rounds of 3D classifications, yielding a dataset with 560,366 particles, generating a 3.3Å-resolution map. Another round of 3D classification focusing the alignment on the complex, resulted in two conformations with high-quality features. After refinement, the resolution levels of these two maps improved to 3.43 Å and 2.99 Å. Particle subtractions on the ECD and TM domains were also performed to further improve the map quality. After several rounds of 3D classifications, ECD map has a resolution of 3.07 Å with 493,869 particles, while that for TM is 4.3 Å with 389,105 particles.

For the CaSR inactive state dataset, the 3D classification resulted in extraction of 55% good particles with a pixel size of 1.071 Å. The particles were subsequently subjected to an auto-refine procedure, yielding a 6.0-Å-resolution map. Afterwards, particles were further sorted another round of 3D classification focusing the alignment on the TM domain, resulted in 37.7% particles with high-quality features. Further 3D classification on the whole complex separates three different orientations of ECD relative to TM domain. After refinement, the resolution levels of these three maps improved to 5.79 Å, 6.88 Å and 7.11 Å. Particle subtractions on the ECD and TM domains were also performed to further improve the map quality. After several rounds of 3D classifications, ECD map has a resolution of 4.5 Å with 253,294 particles, while that for TM is 4.8 Å with 691,246 particles.

### **M**odel building and refinement

The crystal structure of CaSR ECD in apo and active form (PDB Code: 5K5S, 5K5T) were used as initial templates for the ECD of the CaSR. The cryo-EM structures of mGluR5 in resting and active form (PDB Code 6N51, 6N52) were used as initial models for the TM domains of the receptor. The agonist TNCA was generated by COOT (Emsley and Cowtan, 2004) and PHENIX.eLBOW (Adams et al., 2010). The initial templates of ECDs and TMDs were docked into the cryo-EM maps of CaSR using UCSF Chimera (Goddard et al., 2018) to build the initial models of CaSR in inactive and active form. Then the initial models were manually rebuilt the main chains and side chains in COOT. The models were subsequently performed by real-time refinement in PHINEX.

### Intracellular Ca^2+^ flux assay

HEK293T cells were transiently transfected with wild-type or mutant full-length CaSR plasmids. 5μg DNA plasmid was incubated with 15μl lipofectimin in 500μl OptiMEM for 10 minutes at room temperature and then added to the cells for overnight incubation at 37℃. The transfected cells were trypsinized and seeded in 96-well plates. On the day of assay, the cells were incubated with loading medium containing 20mM HEPES, 125mM NaCl, 4mM KCl, 1.25mM CaCl_2_, 1mM MgSO_4_,1mM Na_2_HPO_4_, 0.1%D-glucose and 0.1%BSA at 37℃for 4 hours. Then the buffer was replaced with 100μL of buffer containing Fluo-4 at 37℃ for 1 hour incubation, and then placed into the FLIPR Tetra High Throughput Cellular Screening System. Data was analyzed by non-liner regression in Prism (GraphPad Software). Data points represent average ± s.e.m of quadruplicate measurements.

### Surface plasmon resonance

Surface plasmon resonance (SPR) experiments were performed using a Biacore T2000 instrument (GE Healthcare). The system was flushed with running buffer (20 mM HEPES pH 7.4, 150 mM NaCl, 0.05% Tween 20) and all steps were performed at 25°C chip temperature. The CaSR ECD flowed through the negatively charged chip at a concentration of 1 mg/mL and a flow rate of 10 μL/min for 1 min and was captured by amino-carboxyl coupling reaction. It was followed that the nanobody NB-2D11 went through the chip at a series of concentration (30 μL/min, association: 90 s, dissociation: 220 s). All biacore kinetic experiment data were obtained using Biacore S200 Evaluation Software to calculate the *K*_D_, which is the ratio of *k*d/*k*a.

## Acknowledgements

The cryo-EM data were collected at the Cryo-Electron Microscopy Research Center, Shanghai Institute of Materia Medica, Chinese Academy of Sciences. This work is supported by National Natural Science Foundation of China (No. 31670743), the Strategic Priority Research Program of the Chinese Academy of Sciences (No. XDA12040326), Science and Technology Commission of Shanghai Municipality (No. 3918JC141540001), Joint Research Fund for Overseas, Hong Kong and Macao Scholars (No. 81628013),Natural Science Foundation of Shanghai (16ZR1442900), National Science Foundation for Young Scholar projects (118180359901) and The grand from the Shanghai Institute of Materia Medica, the Chinese Academy of Sciences (CASIMM0120164013, SIMM1606YZZ-06, SIMM1601KF-06, 55201631121116101, 55201631121108000, 5112345601, 2015123456005).

## Author contributions

Y.G. conceived the study. X.C.C. and Y.G. developed purification schemes and purified all proteins for cryo-EM studies; L.W, X.C.C., Z.Y.D and Y.G built and refined models of full-length CaSR from cryo-EM data; Y.G., X.C.C. and L.W. wrote the manuscript; X.C.C. prepared cryo-EM grids and collected cryo-EM data; Y.G., Y.Z.D., L.W and X.C.C. processed cryo-EM data; X.C.C screened conditions for freezing cryo-EM grids of CaSR. Y.J.K., W.Q.Z., H.N.W., X.M.J., M.D. and Z.Z.S. performed llama immunization, cDNA production, and selections by phage display. Y.J.K., L.H., and W.Q.Z. performed early characterizations of CaSR. X.C.C., Q.Q.C. L.H., Y.Y.L., X.Y.L., and M.D performed all in vitro characterizations of Nanobody. Y.G supervised the project.

## Competing interests

The authors declare no competing interests.

## Data and materials availability

All data is available in the main text or the supplementary materials. Cryo-EM maps of active CaSR in complex with TNCA and inactive CaSR in complex with NB-2D11 have been deposited in the Electron Microscopy Data Bank under accession codes: EMD-30997 (NB-2D11 bound CaSR), EMD-30996 (TNCA bound CaSR). Atomic coordinates for the CaSR in complex with TNCA or NB-2D11 have been deposited in the Protein Data Bank under accession codes: 7E6U (NB-2D11 bound CaSR), 7E6T (TNCA bound CaSR).

## Supplemental Information

**Figure 1-figure supplement 1.**
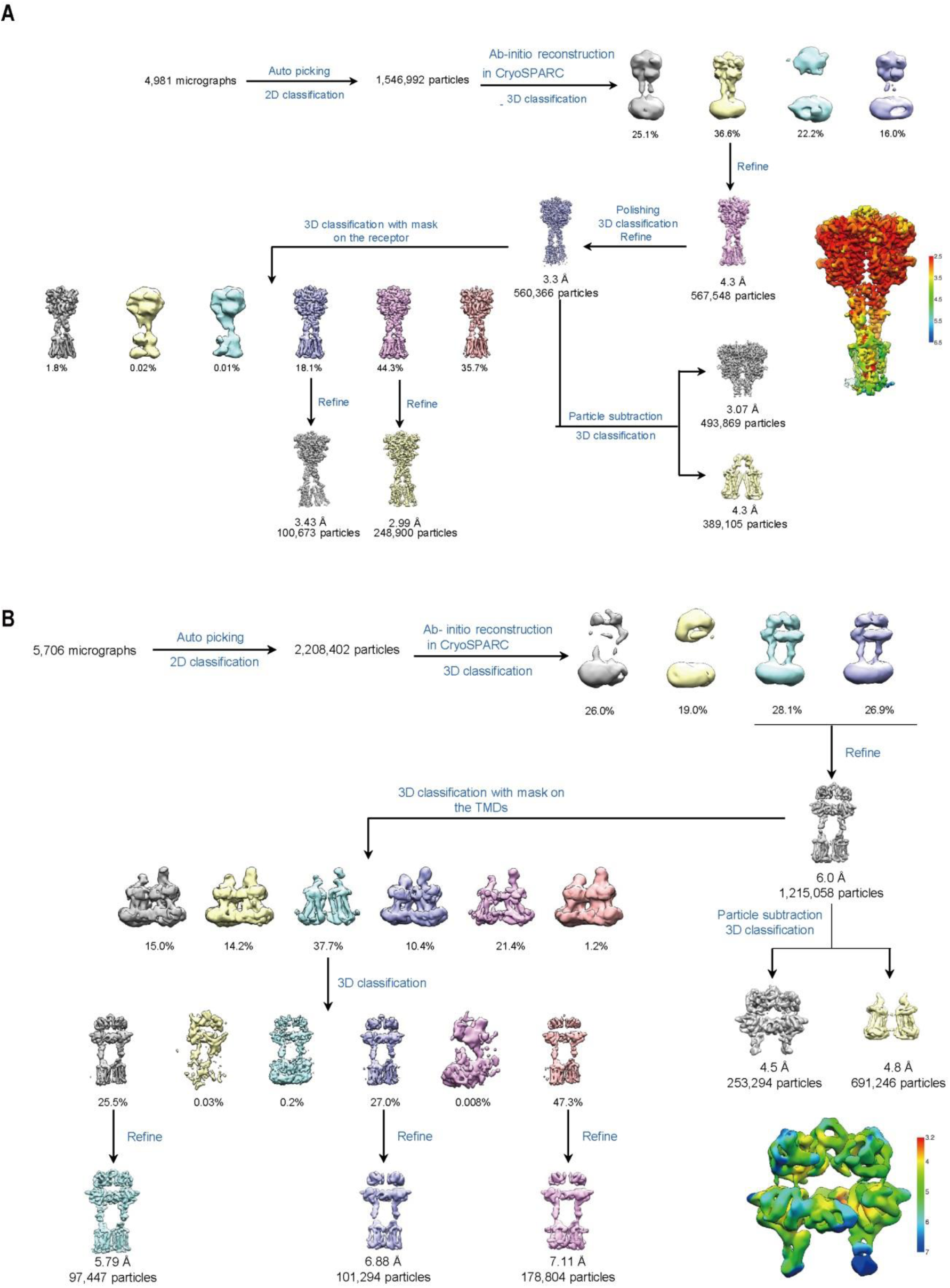
Cryo-EM processing workflow of CaSR. **(A)** The flow chart displaying the cryo-EM processing of active CaSR in GDN, along with the local resolution map. **(B)** The flow chart displaying the cryo-EM processing of inactive CaSR bound with NB-2D11 in GDN, along with the local resolution map.

**Figure 1-figure supplement 2.**
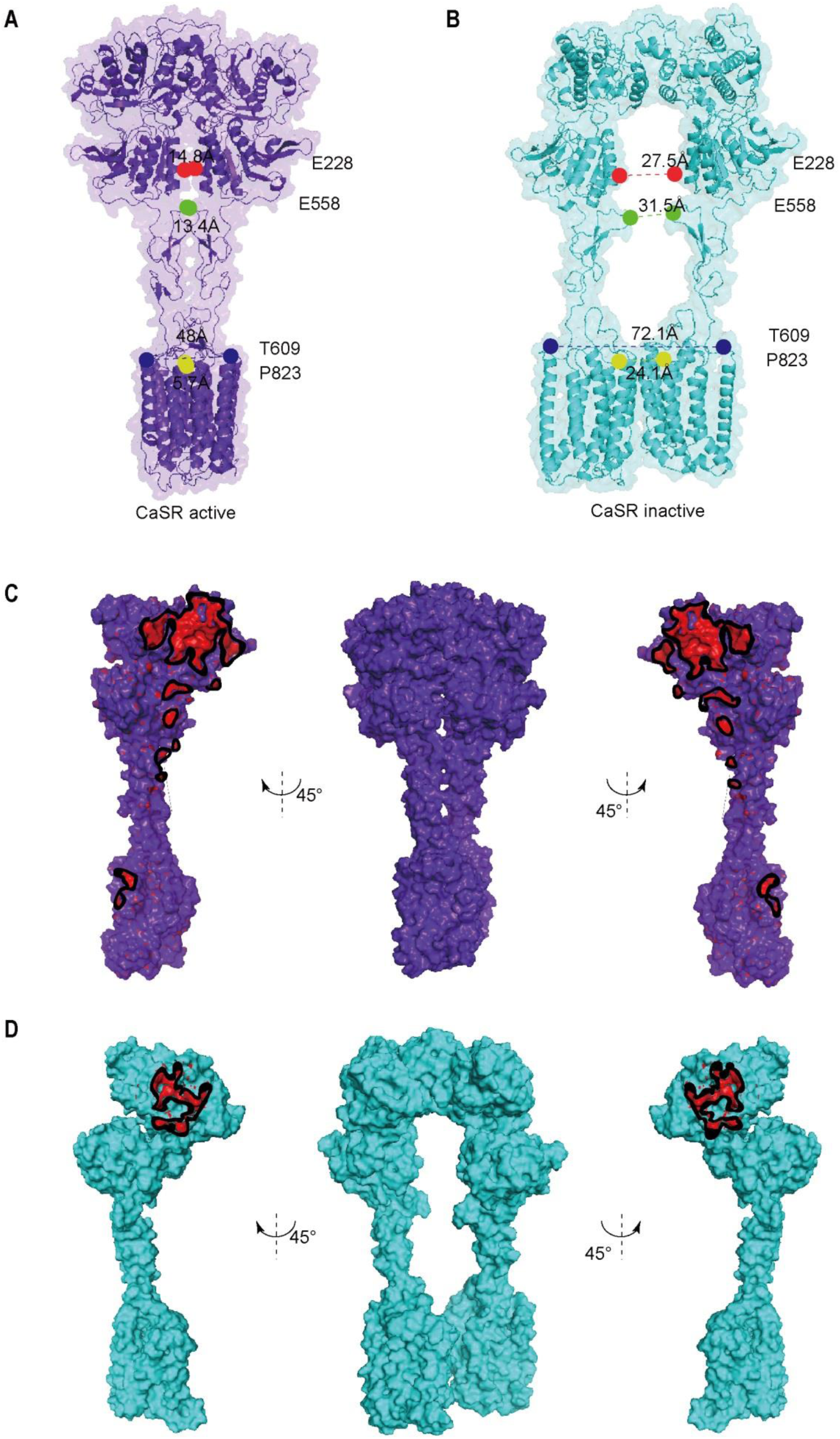
Cryo-EM maps and models of CaSR. **(A-B)** The full-length CaSR cryo-EM maps and models are shown view from front view in the active state (purple) (A), and the inactive state (cyan) deletion of NB-2D11 for clarity (B). Positions in the VFT (red, E228), CRD (green, E558), CRD/7TMD interface (blue, T609) and 7TM domain (yellow, P823) show that the active state is characterized by smaller intersubunit distances. The P823 position in the active model (yellow) shows the contact of 7TMDs. **(C-D)** Comparison of the intersubunit interfaces in active and inactive CaSR are shown for active (C) and inactive (D) CaSR. Contact regions (red) show residues within 4 Å of the opposite subunit. It should be noted that there is only one LB1-LB1interaction interface for the inactive CaSR, and its total buried surface area is much smaller than that of the active CaSR with four different contact interfaces.

**Figure 3-figure supplement 1.**
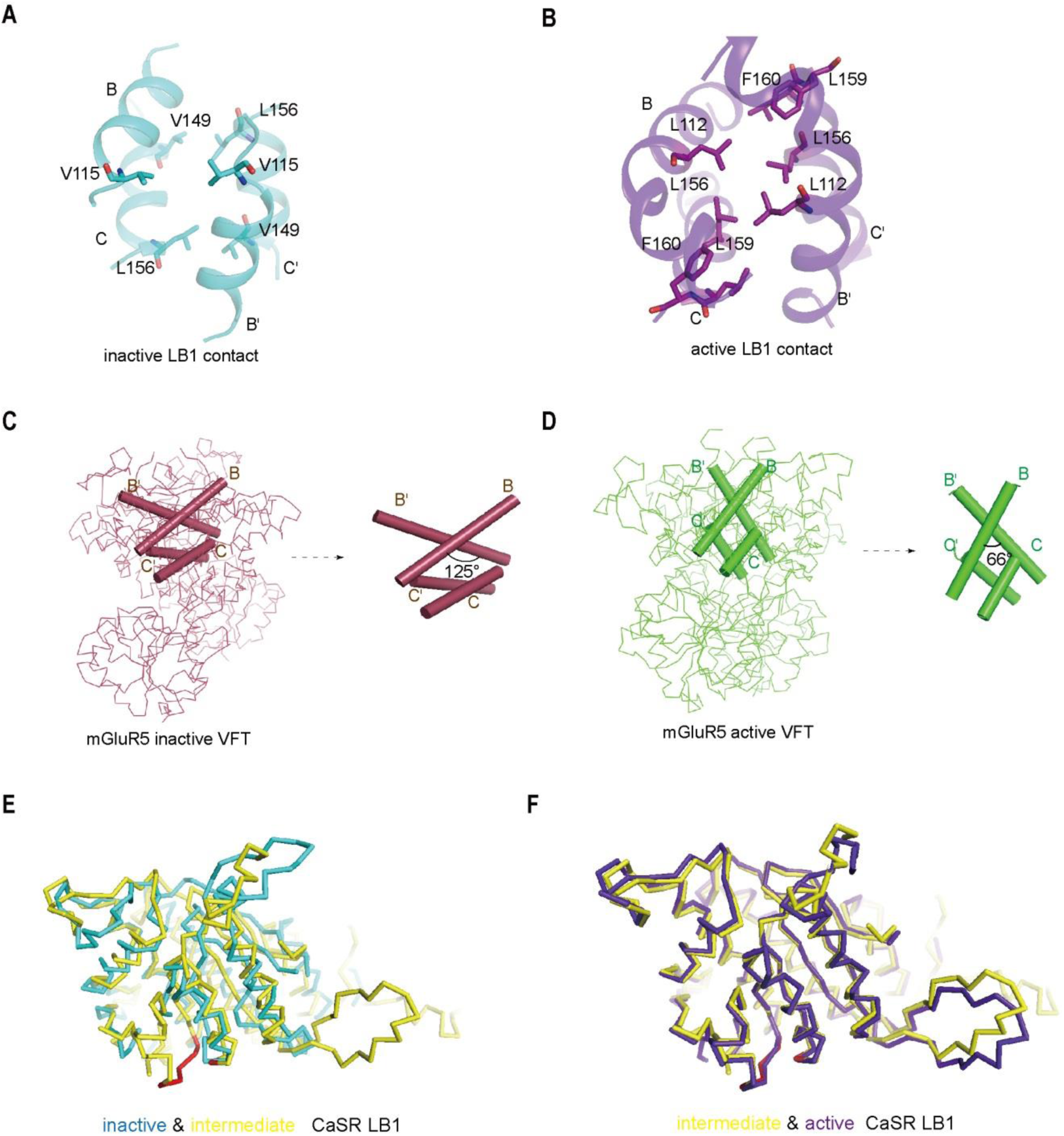
Comparisons of intersubunit LB1 domains interfaces in the inactive and active states of CaSR. **(A-B)** The interface is a hydrophobic patch between residues on the B and C helices of each promoter. In the inactive conformation, it is an interface involving V115, V149 and L156 residues (A, cyan), whereas LB1 interface of the active state (B, purple), is packed with residues L112, L156, L159, and F160. **(C)** Left panel: The Cα trace of VFT module of inactive mGluR cryo-EM structure (raspberry) (PDB: 6N52). Right panel: The B-Helix angle is 125°。 **(D)** Left panel: The Cα trace of VFT module of active mGluR cryo-EM structure (green) (PDB: 6N51). Right panel: The B-Helix angle is 66°。 **(E-F)** Alignment of LB1 domain of CaSR in three conformations: inactive (cyan), intermediate (yellow, PDB:5K5T) and active (purple) states, ligand binding residues (red). (E) inactive and intermediate LB1 domain, the conformation of the ligand binding region in LB1 domain is significantly different in two states. (F) intermediate and active LB1 domain, showing a well superposition.

**Figure 4-figure supplement 1.**
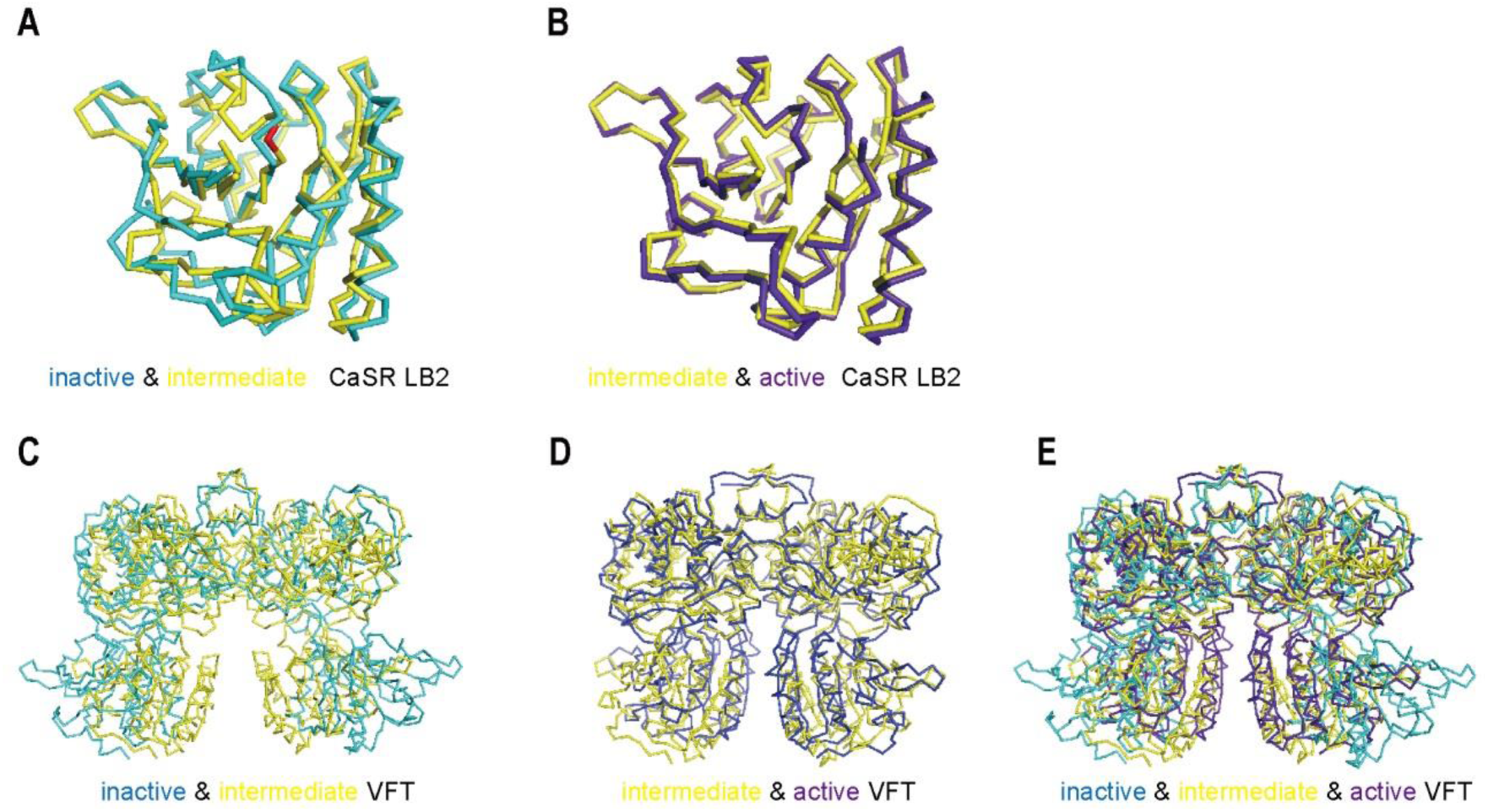
Superposition of LB1 and VFT domains of CaSR. (**A**) inactive and intermediate LB2 domain, **(B)** intermediate (PDB:5K5T) and active LB2 domain, which indicates the three conformations of LB2 domain have no distinguish difference, suggesting the ligand binding residues in LB2 domain do not change during approach of LB2 domain. **(C-E)** Superposition of VFT module based on LB1 domains in the inactive and intermediate (C), intermediate and active (D), and the superposition of inactive, intermediate and active states of CaSR (E).

**Figure 6-figure supplement 1.**
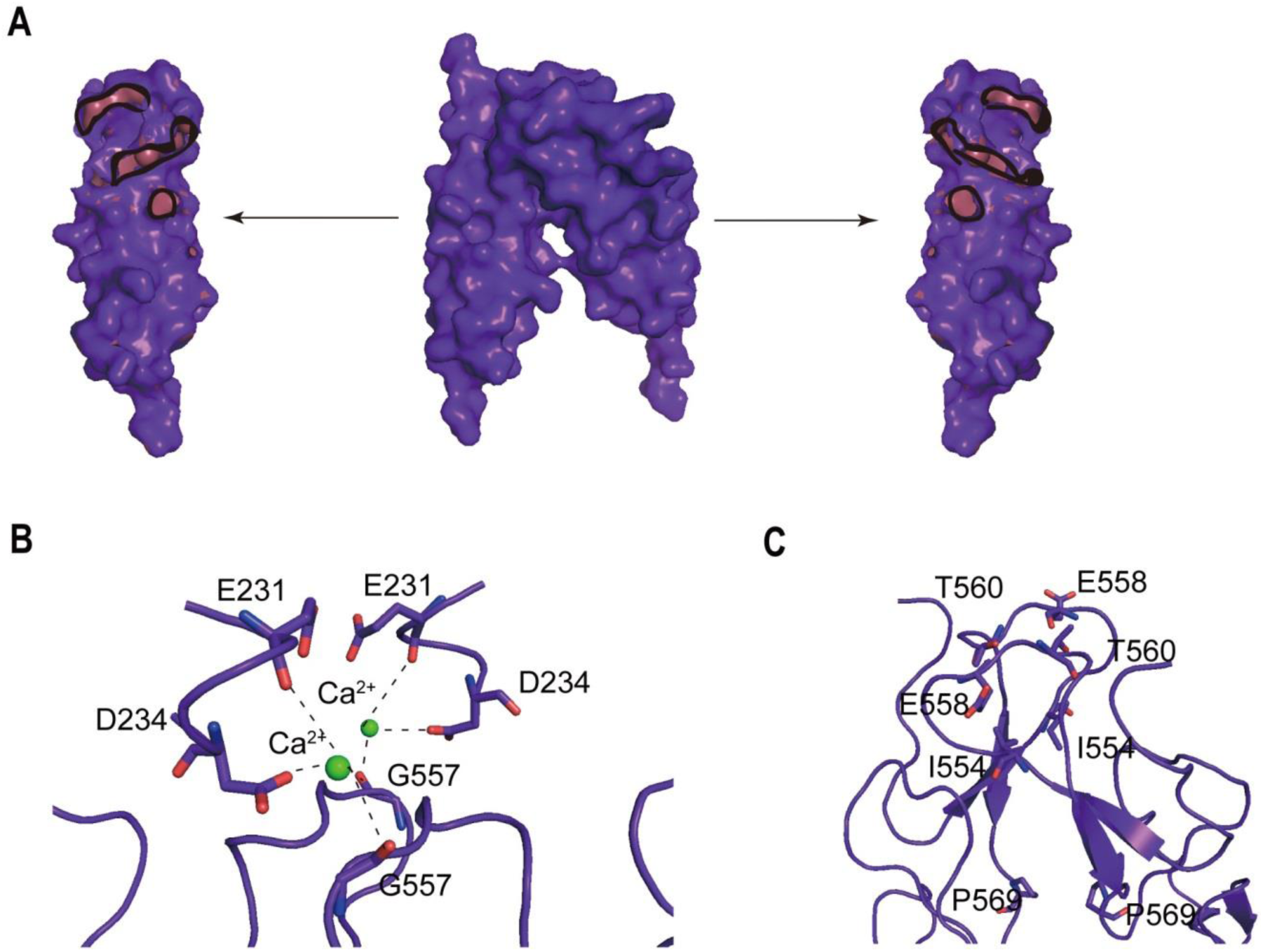
The homodimer interface of the CRDs in the active state of CaSR. **(A)** The homodimer interface of the CRDs (surface represent purple) . Contact regions (red) show residues within 4 Å of the opposite. it covers approximately 1079.09 Å^2^ of solvent accessible surface area. (**B**) There is a bound Ca^2+^ coordinated by carboxylate group of D234 and carbonyl oxygen of E231 and G557, holding the interface that is required to activate the receptor. **(C)** The CR-CR contacts were maintained through the cross-subunit hydrogen bonds between T560 and E558, and hydrophobic interaction of I554 and P569.

**Figure 6-figure supplement 2.**
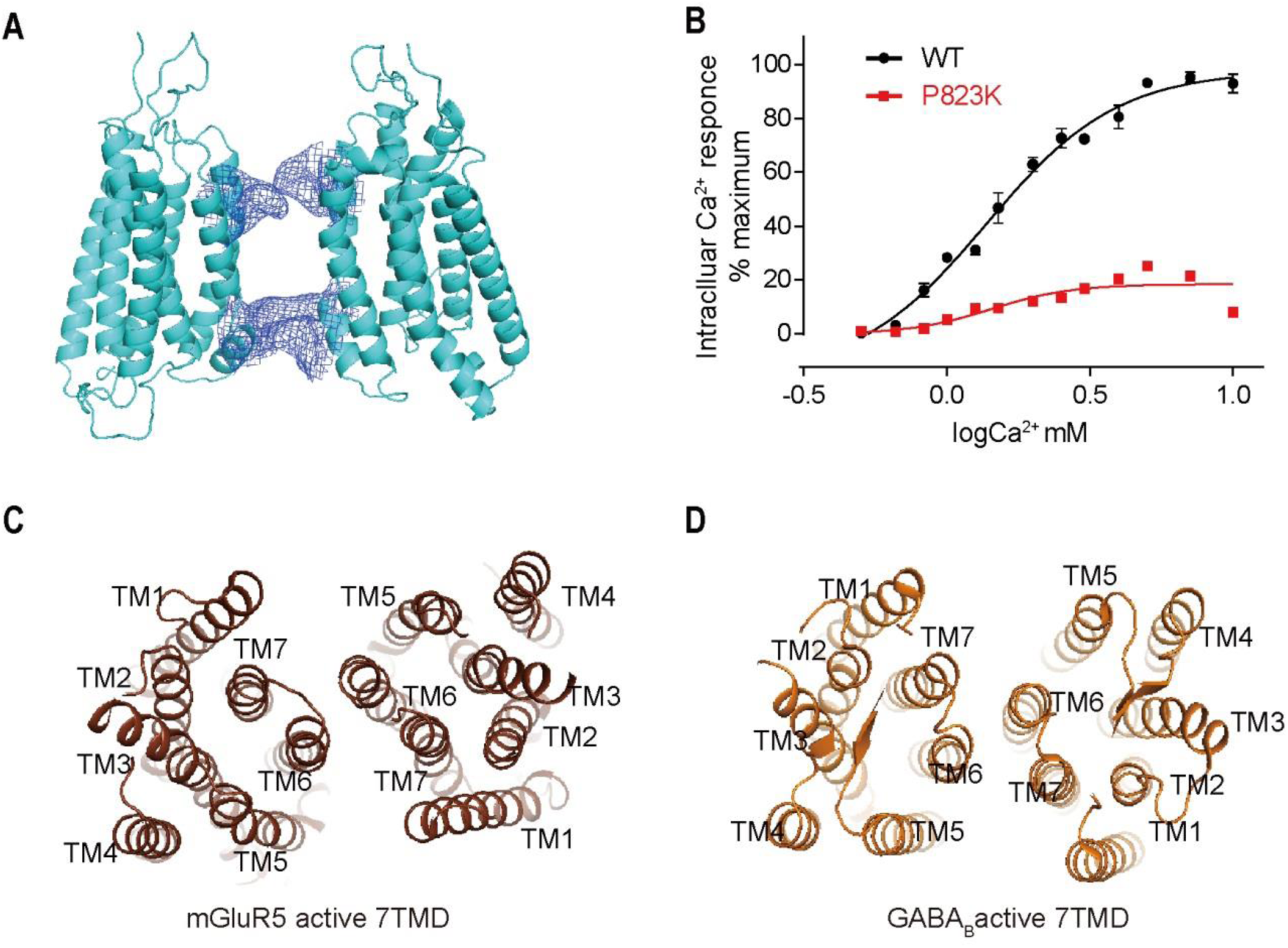
Conformational change of the 7TMDs interface during activation. (**A**) The 7TMD configuration in the inactive state from front view. There are pairwise symmetrical undefined maps that link the extracellular and intracellular part of TM5 and TM6 in the 7TM dimer interface, blocking the association of 7TMDs could regulate the function of CaSR. **(B)** P823K mutant were designed based on structure to disrupt the intersubunit interaction of 7TMD. N=4, Data present mean ± SEM. **(C-D)** The 7TMDs interface of the active state of mGluR5 (C, PDB:6N51) and GABA_B_ (D, PDB:7C7Q) have the interface contacted with TM6.

## List of source data

Figure 2-source data 1 **Intracellular Ca^2+^ flux assay on various CaSR mutations.**

Figure 5-source data 1 **SPR sensorgram of NB-2D11 binding affinity.**

Figure 5-source data 2 **Intracellular Ca^2+^ flux assay on CaSR-NB-2D11 complex.**

Figure 6-source data 1 **Intracellular Ca^2+^ flux assay on various CaSR mutations.**

Figure 7-source data 1 **Intracellular Ca^2+^ flux assay on various CaSR mutations.**

Figure 6-figure supplement 2-source data 1 **Intracellular Ca^2+^ flux assay on CaSR mutation.**

## Notes

### Competing Interest Statement

The authors have declared no competing interest.

